# *Drosophila* antennae are dispensable for gravity orientation

**DOI:** 10.1101/2023.03.08.531317

**Authors:** Nikolay Kladt, Michael B. Reiser

## Abstract

The nearly constant downward force of gravity has powerfully shaped the behaviors of many organisms [1]. Walking flies readily orient against gravity in a behavior termed negative gravitaxis. In *Drosophila* this behavior is studied by observing the position of flies in vials [2–4] or simple mazes [5–9]. These assays have been used to conduct forward-genetic screens [5, 6, 8] and as simple tests of locomotion deficits [10–12]. Despite this long history of investigation, the sensory basis of gravitaxis is largely unknown [1]. Recent studies have implicated the antennae as a major mechanosensory input [3, 4], but many details remain unclear. Fly orientation behavior is expected to depend on the direction and amplitude of the gravitational pull, but little is known about the sensitivity of flies to these features of the environment. Here we directly measure the gravity-dependent orientation behavior of flies walking on an adjustable tilted platform, that is inspired by previous insect studies [13–16]. In this arena, flies can freely orient with respect to gravity. Our findings indicate that flies are exquisitely sensitive to the direction of gravity’s pull. Surprisingly, this orientation behavior does not require antennal mechanosensory input, suggesting that other sensory structures must be involved.

## Results

To investigate the effects of the downward pull of gravity on walking fruit flies, we introduced flies into a tilted arena that was covered by a glass disc to prevent flight (Figure 1A, side view) [17]. The arena was placed within a dark enclosure to ensure that the flies could not use visual cues to guide their paths. By setting the tilt angle of the arena prior to an experiment, we were able to vary the amplitude of the downhill force experienced by walking flies. When flies walk in the center of the arena and away from the walls, the gravitational pull is the dominant directional cue. We implemented a mechanical shutter-like mechanism to relocate flies to the center of the arena (Figure 1A, top view; Figure S1); this enabled repeated measurements of gravity responses while providing a mechanical startle that defined the start of each trial (Figure 1B; Movie S1). Groups of 25 male or female flies were filmed under infrared illumination, and the videos were subsequently tracked [18], to yield trajectories of individual flies. A comparison of example trajectories of flies walking in a horizontal and a vertically tilted arena illustrates the dramatic behavioral changes caused by the substrate incline (Figure 1C). In contrast to the horizontal configuration, flies in a tilted arena walked upwards with straighter paths, and did so from the beginning of the trial (Figure 1C, Movie S1). Flies are indeed startled at the onset of each trial. In both the horizontal and vertical conditions flies initially showed an average speed of about 10-15 mm/s, that declined over the next 10 s to an average speed of less than 5 mm/s (Figure 1D). Furthermore, the flies moved towards the arena boundary within 10 s and remained near the edge throughout the trial (Figure 1D). Flies on a horizontal platform will avoid open spaces and prefer to be near edges [19, 20]; preventing long periods of edge-bound behavior was the primary motivation for developing the shutter mechanism.

**Figure 1:**
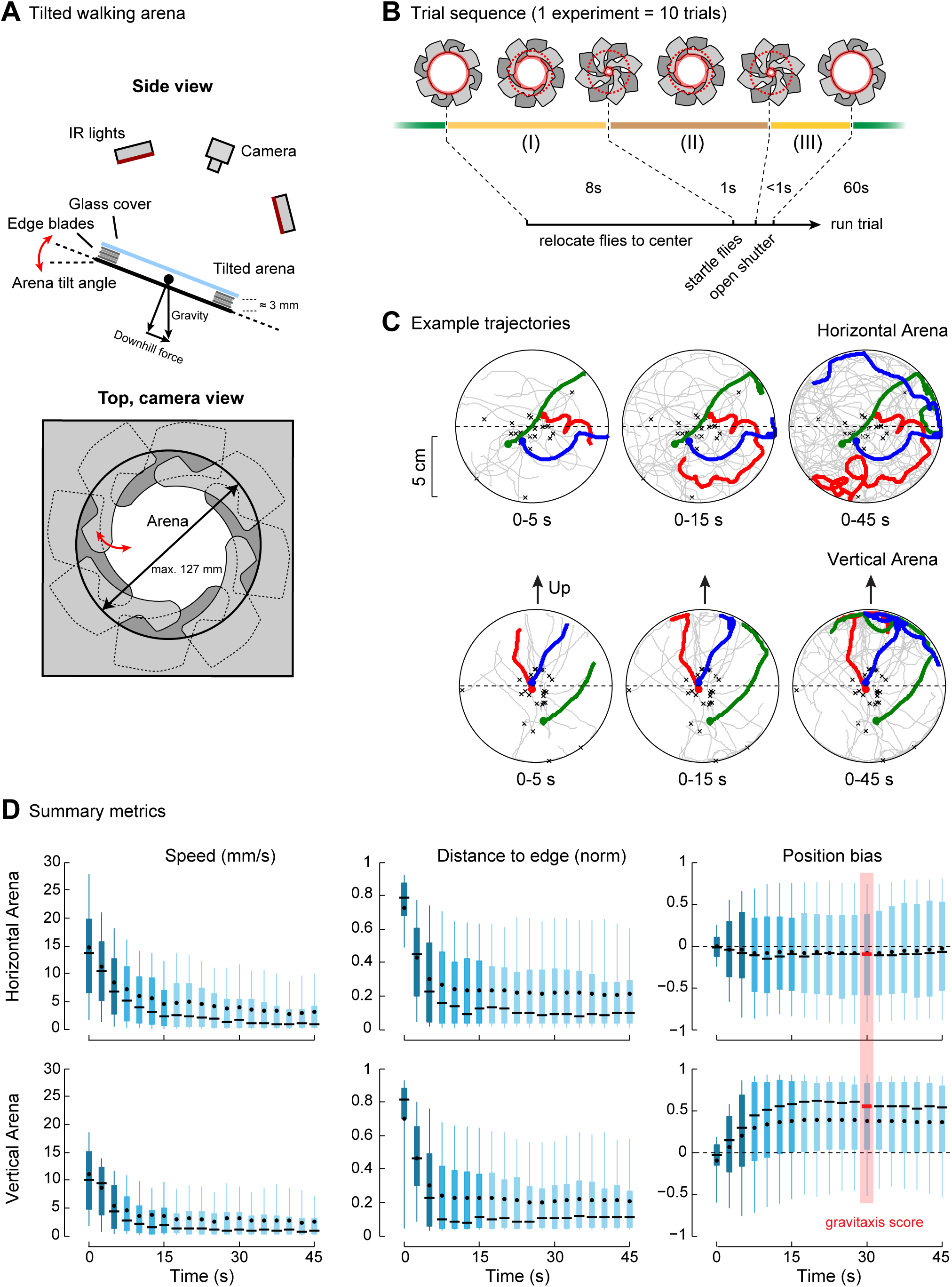
*Drosophila* walk upward in a tilted arena. (A) Side view: the tilt angle of the arena determines the downhill force experienced by walking flies. Flies are placed in the arena; a glass lid encourages walking on the arena floor. Video recordings of flies were captured under infrared illumination. Top, camera view: To relocate flies to the center of the arena, the chamber’s edge was composed of a shutter-like mechanism of interleaved blades that restrict the exposed arena floor by closing. The radius can be adjusted between 15 mm and 63.5 mm. (B) The behavior of groups of 25 flies is recorded over 10, 60 s trials. Prior to the start of each trial, the blades are closed (over 8s) to gently push flies to the center of the arena. A rapid open-close sequence (1s) is used to startle the flies, followed by opening the walls to the maximum radius (<1s), which defines the start of each trial. In off-line tracking of flies, the trajectories are initialized once the shutter is completely open. (C) 30 randomly chosen trajectories from a single experiment (in gray) demonstrate the differences between locomotion in the horizontal and vertical arena. Three example trajectories are highlighted as red, green and blue; the initial position of each trajectory is marked with a black ‘x’. The development of the trajectories over time is shown by plotting these trajectories from the start of the trial until 5, 15, and 45 seconds. In the tilted arena, flies move upwards, as most of the trajectories are above the dashed line. (D) The distributions of summary statistics are obtained by pooling the speed, absolute distance to the arena edge, and position bias, across flies and trials for the horizontal (N=259 trajectories) and vertical arena condition (N=286 trajectories) of a single experiment. The position bias measures the normalized height of the flies along the tilt axis, where a 1 indicates that all flies are at the top. Boxplots present the data in 3 s intervals, indicating the median (black bars), the mean (black dots), the lower and upper quartiles (boxes), and the 0.1 and 0.9 quantiles (vertical lines). The gravitaxis score (red bar) is defined as the median position bias for the 30 s bin.

To examine the gravity-specific behavioral responses, we define the *position bias* of the flies to be the ‘vertical’ height relative to the boundaries of the arena (see Materials and Methods). This metric confirms what is plainly visible in the example trajectories (Figure 1C)—flies in the vertical arena walked upwards, whereas flies in the horizontal chamber did not show a directional preference and distributed uniformly around the arena’s midline (Figure 1D). The average fly position in the arena shows a strong bias towards the upper edge of the arena throughout the trial (Figure 1D). In both the horizontal and vertical arena the distributions of these metrics are rather stable from ∼20 seconds onwards. We therefore define the median position bias at 30 seconds to be the ‘gravitaxis score’ (Figure 1D), since it captures the consequence of gravity-based orientation (see Materials and Methods).

To test the sensitivity of walking *Drosophila* to a range of downhill forces, we examined the behavioral responses of flies confronting different tilt angles (Figure 2A). As expected, the gravitaxis score for the horizontal condition is near zero. To our surprise, flies exhibited a significant gravitaxis score at the 5° tilt (p<0.05), and the gravitaxis score is approximately constant for arena tilts between 20° and 90° (Figure 2A; p<0.01; results further analyzed in Figure S2A-C).

**Figure 2:**
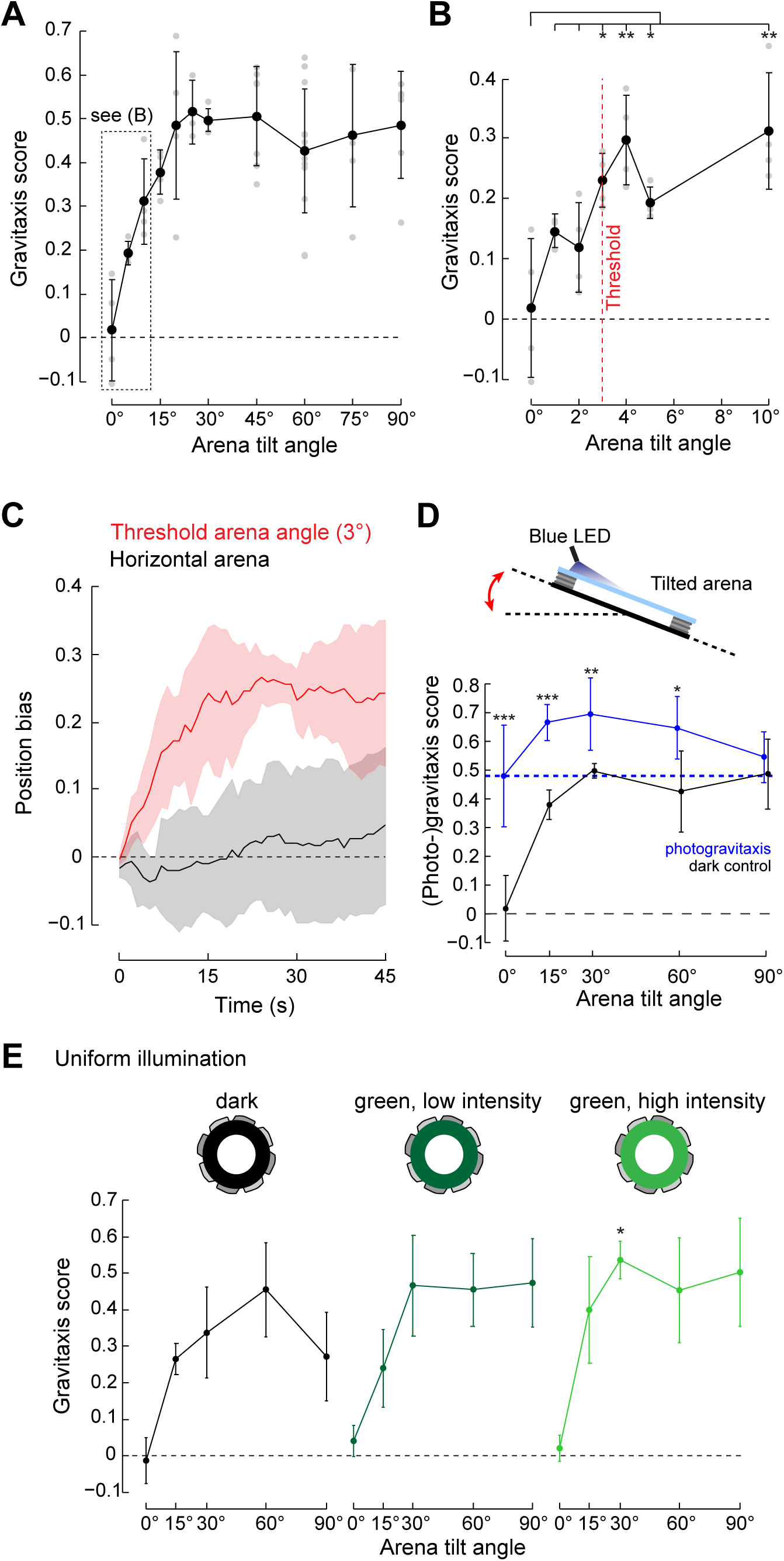
Flies exhibit gravitaxis on small inclines. (A) The mean gravitaxis scores are plotted for a range of tilt angles (black dots are mean ± s.d., gray dots are the results of individual experiments, N ≥ 4 experiments per tilt angle). The gravitaxis score for all tilt angles is significantly different from the horizontal condition (p<0.05 for 5°, p<0.01 for 10°, 15°, 20°, 25°, 75°, p<0.001 for 30°, 45°, 60°, 90°; two-tailed t-test). (B) The minimum detectable incline was determined by testing arena tilt angles between horizontal and 5° in 1° steps. Flies walking on tilt angles of 3° and higher show a gravitaxis score that is significantly larger than for the horizontal condition (two-tailed t-test). (C) Flies walking in the arena with a 3° tilt show a rapid upward response, indicated by a positive position bias. This bias was maintained over time (mean ± s. d., N=4 experiments). (D) Phototaxis was tested by fitting the arena with a blue LED (470 nm), lighting the top edge of the tilted arena. Dark control data are a subset of the results presented in Figure 2A. At most tilt angles, the combination of gravitaxis and phototaxis results in greater upwards movement as compared with the gravitaxis alone response (N = 11 experiments for 0°, N = 4-5 experiments for 15°, 30°, 60°, 90°. P-value/Arena tilt: 0.0004/0°, 0.0004/15°, 0.008/30°, 0.012/60°, 0.43/90°). (E) The effect of uniform surround illumination was tested using a controllable green (568 nm) LED display. Two different green intensities were tested and compared to dark control experiments conducted in the same arena. Each condition (lighting, incline combination) was tested 4 times. The gravitaxis scores are not significantly different at each arena tilt angle for the different illumination conditions, except for the comparison between the dark and high intensity lighting at 30° tilt (p=0.027, two-tailed t-test).

To determine the smallest incline that flies detect, we tested tilts between 0°-5° in 1° steps (Figure 2B). Although the mean gravitaxis scores for all angles above 0° are positive, the smallest tilt angle for which this difference is statistically significant is 3°, representing a level of sensitivity which, to our knowledge, is much lower than what has been previously reported for fly gravitaxis [21]. Gravity orientation in flies need not be supported by a single sensory structure, and therefore it might be expected that the orientation behavior on small inclines would be rather different than on large tilts. However, there are no qualitative differences between the growth of the position bias over time in the vertical arena (Figure 1D) and this response in the 3° tilted arena (Figure 2C), which both showed an increase in the position bias within the first 10 – 15 s that was maintained throughout the remainder of the trial. The initial response of flies further confirms that flies show a rapid and sustained upward response with a detection threshold of not more than 3° (Figure S2D-F). We also used a simpler assay to independently confirm the general shape of the response curve. In this assay, individual flies emerged through a hole in the center of the arena and we recorded only one response per fly (Figure S2G-I). We note the remarkable agreement between the behavioral responses measured in both assays. Together, these results clearly indicate that the gravitaxis response of *Drosophila* is exquisitely sensitive to even small inclines, and that flies show immediate, upward-directed orientation as soon as they are confronted with an inclined surface. Establishing the sensitivity of gravity orientation is critical for determining the necessary capabilities of the underlying sensory structures that support this behavior.

Would access to additional sensory information influence the upward movement of flies on inclined surfaces? All experiments reported thus far were performed in the dark so that flies could not use any visual information for orientation. Phototaxis behavior, in which many seeing organisms orient towards a source of light [22], is known to produce strong orienting responses in *Drosophila* [23]. We tested combinations of gravitaxis and phototaxis by placing a blue LED at the perimeter of the arena [24], while measuring behavior at 5 different arena tilt angles (Figure 2D, see Materials and Methods). Comparing the strength of the position bias (at 30s) for phototaxis in the horizontal arena (dashed blue line) with gravitaxis in the vertical (dark) arena, we see that these two stimuli are equally effective at inducing directed movement of flies (p=0.94). This indicates that under our experimental conditions, these two cues independently lead to similarly strong taxis. When presented together, we see that there is a modest, but significant enhancement of directed displacement of flies, especially at the lower tilt angles (Figure 2D).

Would visual information that does not provide a directional cue, such as access to the spatial layout of the chamber, also enhance the gravity response? To test this possibility, we configured our experiment to provide uniform illumination from the perimeter of the chamber. Three different conditions were tested: dark, low and high intensity green lighting (see Materials and Methods) for 5 arena tilt angles (Figure 2E). In the horizontal arena configurations, there is no directional bias due to the surrounding illumination (Figure 2E, 0°), while the gravitaxis scores at other tilt angles are not significantly different between the three lighting conditions (with one exception; Figure 2E). Taken together, these results suggest that the robust orientation upwards on inclined surfaces is neither impaired nor enhanced by the addition of non-directional visual information.

We have shown that flies vigorously move upwards once they encounter an inclined surface, and that this response is strong and rapid even for very small tilt angles. But how do flies achieve this robust upward navigation? To elucidate the fine-scale orientation of flies walking on sloped surfaces without visual cues, we further analyzed the trajectories obtained for the range of arena tilts presented in Figure 2A, B. To capture the turning behavior of flies in response to all possible orientations with respect to the upward direction, we compared the initial and final orientations of all fly trajectories in 15 mm segments (Figure 3A; details in Materials and Methods). This analysis shows that the turning response depends on the fly orientation (Figure 3B). In the vertical arena, the distribution of initial segment orientations shows that most flies start the trial aligned with, or near, the upward direction (Figure 3B, Frequency). Once flies are oriented upwards, they maintain this orientation (mean turning angle of ∼0°). Any deviation away from this ‘preferred’ upward orientation leads to compensatory turning responses (Figure 3B, Turning angle). The sign of the mean turning responses indicates that when flies are initially misaligned, they subsequently turned upwards. The magnitude of these compensatory turns depends on the segment path length used in this analysis, although the nearly sinusoidal relationship between initial fly orientation and turning response is seen over multiple length scales (Figure S3A). The variance in the turning response across orientations is smallest for flies initially facing upwards, indicating that these flies turned less (Figure 3B, initial segment orientation around 0°). The decreased variance in upward oriented flies (and increased variance of downward oriented flies) is captured by the mean response vector strength (Figure 3B, R; see Materials and Methods). A summary of turning responses as a function of the initial segment orientations for all tested arena tilt angles (Figure 3C) is in agreement with the gravitaxis score results of Figure 2A—that is flies responded to small tilt angles with a gravity response that is very similar across angles. By examining the turning response of sideways oriented flies over a range of segment lengths, we see a rather consistent, approximately linear trend across arena angles greater than 5° (Figure S3B), suggesting that flies continued to reorient upwards over several length scales. Taken together, these findings suggest that in order to move up inclined surfaces, flies employ the following strategy: (1) when not aligned with the upward direction, they turn so as to minimize this misalignment with a turn amplitude that depends on the degree of misalignment (Figure 3B), and (2) when oriented upwards they suppress all turning as they continue to walk upwards (Figure 3D).

**Figure 3:**
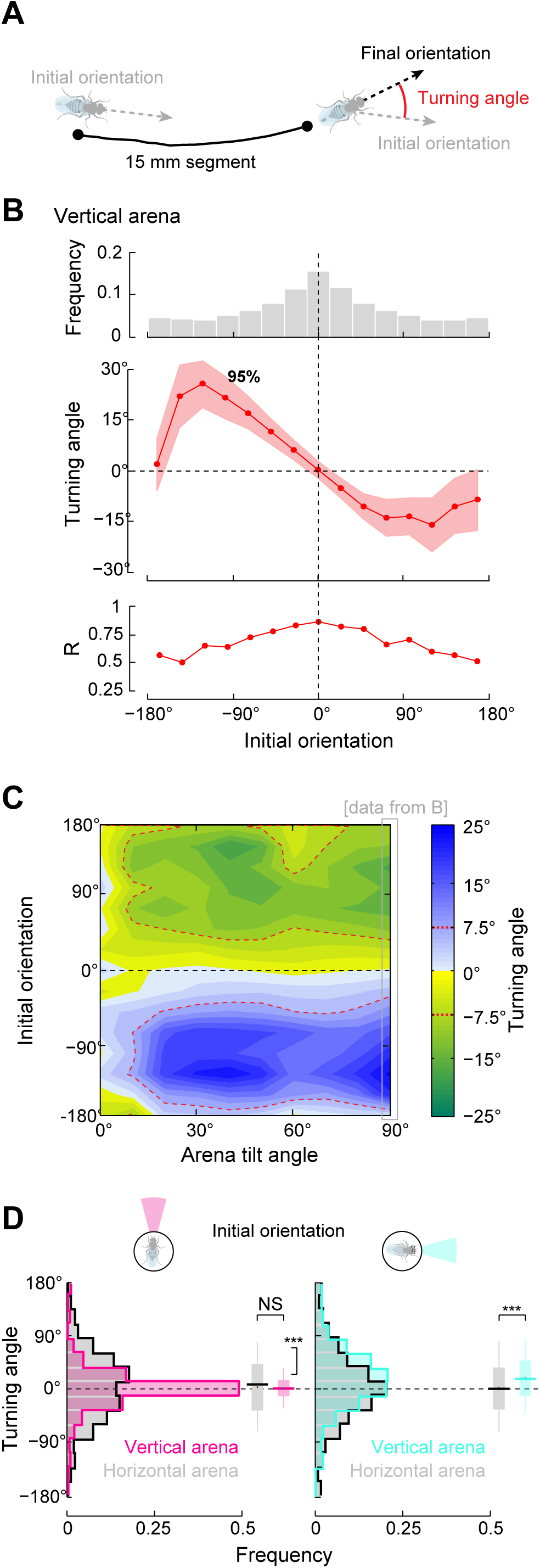
Directed upward walking results from changes in turning behavior. (A) The turning behavior of flies was measured as the difference between the initial and final fly orientation in 15 mm segments. (B) The distribution of all segments (binned by their initial orientation, 24° bins) for the vertically tilted arena (6514 segments from 6 experiments). The plot shows the relative number of segments in each bin (Frequency), circular mean turning angle, and the mean response vector length R. The red dots represent the mean turning angle, shaded red band denotes the 95% confidence interval. (C) The turning behavior depends on both fly orientation and the arena tilt angle. The analysis method used in B is applied to the trajectory segments of flies for all tested tilt angles and summarized using false color. To aid in the visualization of these data, we use a turn angle threshold of 7.5° (red dotted lines), to delineate the rather consistent and directionally symmetric distribution of turning responses of flies that were misaligned with the up direction. The black dashed line shows the upward direction. Results are based on the data from the experiments of Figure 2 (104,883 segments from 13 arena tilt angles). (D) The reactions of flies that are initially oriented upwards (pink) and sideways (light blue; 24° bins; the symmetrical sideways bins were mirrored and data were pooled). Comparing the distribution of turning angles for the vertical arena to the horizontal condition (gray distribution), shows a significant change in the distribution variance of the upwards oriented flies (F-Test, p<0.001; N = 253/581 segments in vertical/horizontal conditions), indicating that flies already walking upwards on an inclined surface suppress their turning. Furthermore, the sideways oriented flies show a significant shift in their mean turning response (Mann-Whitney U-test, p<0.001; N=508/379 segments in vertical/horizontal conditions), demonstrating that they turn upwards.

We next considered possible sensory pathways that might underlie the robust gravity orientation behavior. In flies, several mechanosensory structures have been suggested to play a role in gravity sensing [5, 7, 14, 16, 21, 25]. The antennae have also been implicated in gravity sensing [3, 4, 6, 16, 21, 26] and recent work identified subgroups C and E of antennal Johnston’s organ (JO) neurons as required for normal gravitaxis [3, 4]. We first examined the role of sensory structures on the wings, halteres, and neck and found that the gravitaxis score of flies was unaffected by these surgical treatments (Figure 4A; arena tilt angle of 60°, see Materials and Methods). JO neurons are thought to be activated by torque between antennal segments a2 and a3 that is generated by deflections of a3 and the arista (Figure 4A, antenna schematic). For gravity detection, it is proposed that a static deflection of a3 leads to the activation of highly sensitive C and E neurons [3, 26]. We sought to remove this antennal input by bilaterally ablating a3 and the aristae or by immobilizing the a2-a3 joint (see Materials and Methods). The gravitaxis score for both manipulations is significantly reduced from control levels (Figure 4A), and is unlikely to be caused by surgical manipulations of the flies (since the three previous manipulations were performed under similar conditions). The unilateral ablation of one antenna also resulted in a reduced gravitaxis score (Figure 4A; left a3, arista ablated). We further tested the possible role of the JO C and E neurons by using the GAL4/UAS expression system. We conditionally silenced these neurons by targeting expression of tetanus toxin (TNT) and we limited this expression to adult animals by using a temperature-sensitive GAL80. The results of these genetic manipulations confirm previous findings, since flies reared at 30° C (expressing TNT in JO-CE neurons) show a reduced gravitaxis score in comparison to the identical flies reared at 18° C (blocking expression of TNT; Figure 4A). As these results support the idea that the antennae provide the major sensory input for gravitaxis, it might be expected that flies with impaired antennal function would either be incapable of orienting upwards for all angles or would show a markedly less sensitive response at lower angles. To test this we challenged flies with either the surgical or genetic manipulation of JO function to walk at 5 different arena tilts (Figure 4B). The results of these experiments do not support either expectation. The gravitaxis scores are not different at the lower angles tested, and the antennal effect is only seen at angles greater than 30°. These differences are consistent across trials (Figure S4A). By illuminating the top edge of the arena with a blue LED (as in Figure 2D), we see that these flies are not generally impaired in oriented walking behaviors since phototaxis enhanced the (60°) upward response, restoring it to control levels in antennae ablated flies and slightly improving scores in flies with inactivated JO-CE neurons (Figure 4B, gray dots).

**Figure 4:**
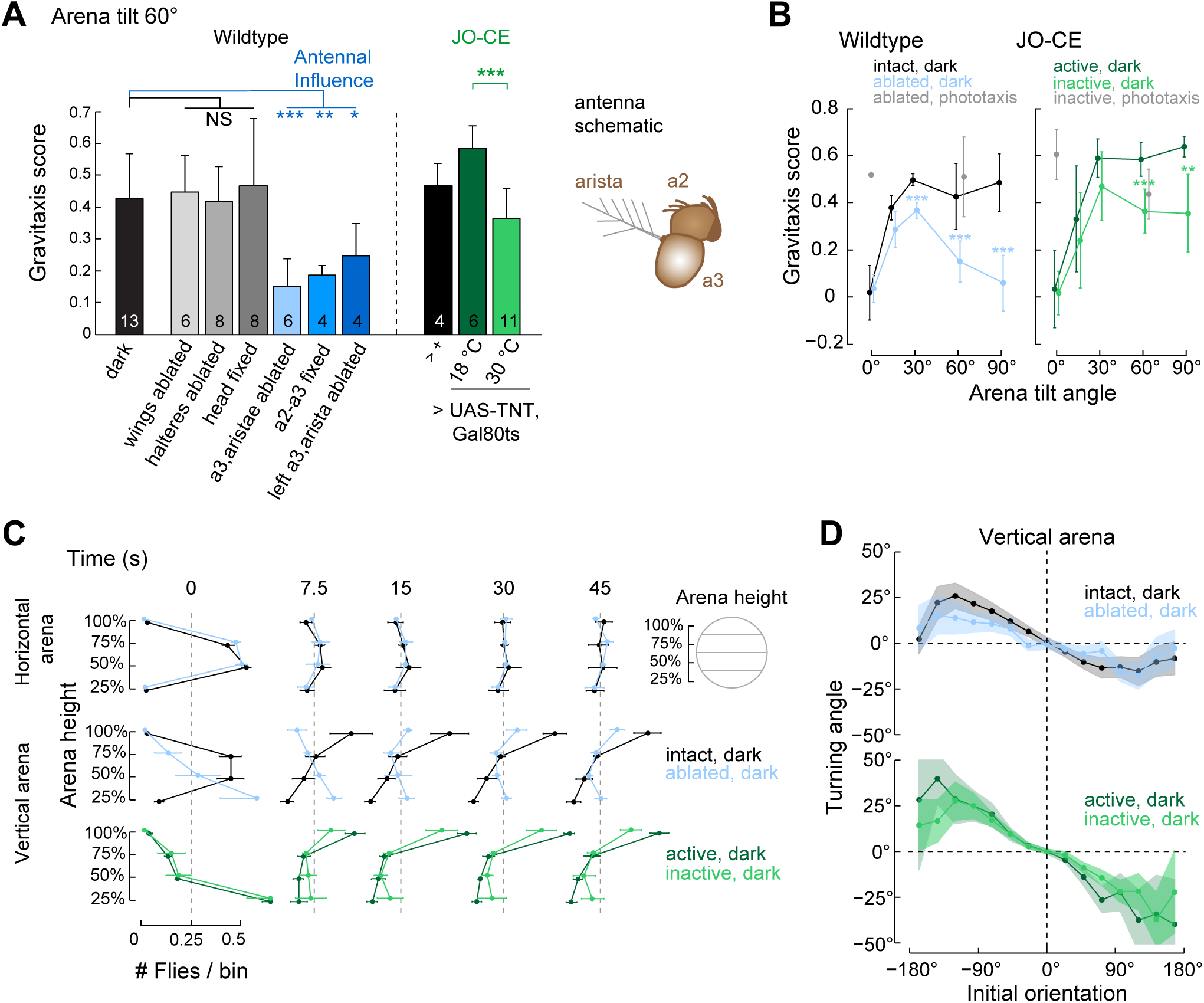
The antennae play a limited role in gravitaxis. (A) The effects of manipulations to several sensory structures were tested in 60°-tilted arena. Surgical ablation of the wings and halters, and fixation of the head, did not reduce the gravitaxis score. Antennal ablation (both uni- and bi-lateral) and fixation resulted in significantly reduced gravitaxis scores. Conditional expression of tetanus toxin (TNT) was used to inactivate the Johnston’s organ (JO) neurons of subclass C and E, which also resulted in a reduced gravitaxis score. The mean gravitaxis scores ± s. d are shown; numbers on bars indicate the number of experiments, all comparisons were performed with a two-tailed t-test (* p<0.05, ** p< 0.01, *** p < 0.001). Antenna schematic indicates the targets of manipulations (see text). (B) The gravitaxis response curves (mean ± s.d.) for bilateral antennal ablations (a3, aristae) and inactivation of JO-CE neurons at five arena tilt angles. In grey, the results of phototaxis trials for horizontal and 60° tilt are plotted (N ≥ 4 experiments / condition). (C) To reveal the antennal effect on gravitaxis over the course of a trial, we quantify the relative number of flies in 4 bins along the height of the arena. The spatial distributions of flies at 5 representative times during the trial are shown. Wildtype flies started in the center of the arena and then distributed uniformly in the horizontal arena, while in the vertical arena they showed a strong upward position bias. In contrast, antennal ablations caused flies to begin trials in the lower part of the arena. By 15 seconds, many of the flies that began at the bottom of the arena moved to the top half. In the JO-CE expressing Gal4-driver line, flies from both conditions began the trial with lower starting positions. By 45 seconds, the control JO-CE flies developed a distribution similar to wildtype control flies (compare the black curve, middle with the dark green curve, lower). Flies with inactivated JO-CE neurons exhibited a bimodal distribution, with more flies at the top of the arena than the bottom (N ≥ 4 experiments / condition). (D) Antennal influence on fly turning behavior is analyzed as in Figure 3B. The influence of the initial orientation on the subsequent turning angle is quite similar between intact and antennae ablated flies and between flies with active and inactive JO-CE neurons. Results based on 90° data from B (N = 6514/intact, 7994/ablated, 4892/active, 10130/inactive, segments per treatment group).

While conducting these experiments, we noticed that, remarkably, the flies with manipulated antennae were often falling to the bottom of the arena after being startled (Movie S2). A comparison of the location of tracked flies along the height of the arena over time confirms this observation (Figure 4C). Wildtype flies began each trial in the center of the arena, and then spread out to cover the entire chamber in the horizontal condition, whereas they moved upwards in the vertical arena. In contrast, most of the antennae ablated flies (Figure 4C; light blue) began in the lower quarter of the vertical (but not the horizontal) arena, but then moved upwards over the course of a trial. This difference is smaller for the comparison of the JO-CE flies reared under the 2 temperature conditions, since both groups exhibited the ‘falling down’ phenotype at the start of the trial, however the difference between the final upward progress of these flies is reduced for the flies with inactivated JO-CE neurons. Since both treatments of antennal function yielded animals that exhibited upward progress on the inclined surface, we further analyzed the turning behavior as a function of initial segment orientation and arena tilt angle (Figure 4D, Figure S4B-C). For both antennal manipulations, the suppression of turning for upward walking flies is reduced (Figure S4B). However, these results indicate that under both conditions, flies with impaired antennal function oriented upwards by turning as a function of their current orientation, and did so with a turning response that is remarkably similar to wildtype flies (Figure 4D, Figure S4B-C).

## Discussion

*Drosophila* gravitaxis has served as an important behavior for measurements of fly locomotion, and is typically studied in various climbing assays [2–4, 11] or gravitaxis mazes [5–8]. The tilted platform we developed combines the quantitative aspects of these approaches with the qualitative features of tilt platform experiments in other fly [21] and insect species [13, 15]. Our approach enables walking flies to freely orient within an arena that can be tilted over a range of angles (Figure 1). Video recording and the use of modern tracking methods [18] enable a rich description of this behavior (Movie S1).

With our setup, we found that flies detect surface tilts of 3° (Figure 2B and S2E) and that the gravitaxis behavior exhibited a maximum response to tilts between 20° and 90° (Figure 2A and S2). The fine-scale analysis of fly trajectories revealed two behavioral modifications that explain the gravitaxis observed on inclined surfaces: flies reduce their misalignment with the up direction by turning with an orientation-dependent amplitude (Figure 3B, C and S3) and upward oriented flies suppress turning and continue to walk straight (Figure 3D). Based on these results, the sensory apparatus that detects the orientation discrepancy must perform over a very large range of body orientations with respect to the environment.

The antennae are the sensory structures that have consistently been implicated as an input to gravitaxis in flies [3-5, 16, 21, 27]. Furthermore, estimates of antennal displacement due to gravity showed that JO neurons have the expected sensitivity to support this function [3], although our evidence for a gravitaxis response at very small tilt angles and the turning dependency on fly orientations dramatically lowers this required sensitivity (Figure 2B, C). Therefore, we tested antennal contribution to gravitaxis with surgical ablation, fixation of the critical antennal joint, and genetic inactivation (Figure 4). Surprisingly, these experiments showed that impaired antennal function led to a reduction of the gravitaxis response at large tilt angles but did not impact the response at lower tilt angles (Figure 4A, B). Although strong reductions in upward movement of flies were seen at larger tilt angles, the turning behavior was nearly indistinguishable between intact and antennae manipulated flies (Figure 4D and S4). This apparent discrepancy is explained by the fact that flies with antennal impairment were likely to fall to the bottom of the arena immediately after the startle that begins each trial (Figure 4C, Movie S2), but once they resumed walking, they exhibited intact gravity orientation. Taken together, these results demonstrate that the antennae cannot be the critical input for gravity orientation. Our suggestion of a limited role for the antennae in *Drosophila* gravitaxis is in agreement with observations in larger flies. For example, in *Calliphora*, removing the antennae increased behavioral variance, while the upward walking bias remained intact. ([21]; an effect that is also captured by our experiments, Figure S4B).

The primary antennal deficit we observed was that flies fell to the bottom of the arena after they were pushed to the center of the arena. This phenotype together with results showing that the antennae are involved in a wind startle induced behavior [28], suggest a specific role for the antennae in mediating reactions to sudden mechanosensory stimuli, that have been collectively referred to as ‘startle responses.’ This proposal is further supported by antennal nerve recordings in flies that showed phasic activity in response to body rotations [4].

In this study, we used a detailed analysis of the behavioral reactions of individual flies to demonstrate that the antennae cannot be the primary sensory input used by *Drosophila* for gravitaxis. Based on previous insect studies, sense organs necessary for graviception are likely located on the insect’s legs [14, 29]. Further studies into the contributions of leg mechanosensors and their interaction with other sensory structures will require even finer analysis of the detailed kinematics of the leg, head, body, and antennae of flies walking on inclined surfaces, which has been made possible by recently developed methods [30]. Mechanosensation in flies has not received the same attention as other sensory modalities such as vision and olfaction, but it will not be possible to truly disentangle any sensorimotor behaviors without a deeper understanding of the natural mechanosensory experience of flies. The integration of sensory cues such as gravity’s downward pull into robust navigation behaviors must have played a crucial role in enabling flies to thrive in diverse environments.

## Acknowledgments

T. Laverty and the Janelia Fly Core assisted with *Drosophila* maintenance. Tanya Tabachnik (Janelia Instrument Design and Fabrication) assisted with the design and assembly of the shutter. Further technical support was provided by M. Bolstad, C. Werner, J. Osborne, and D. Olbris. We thank members of the Reiser and Jayaraman labs, and Janelia Colleagues for providing constant feedback and a stimulating scientific environment. This project was supported by HHMI.

---

**Figure S1.**
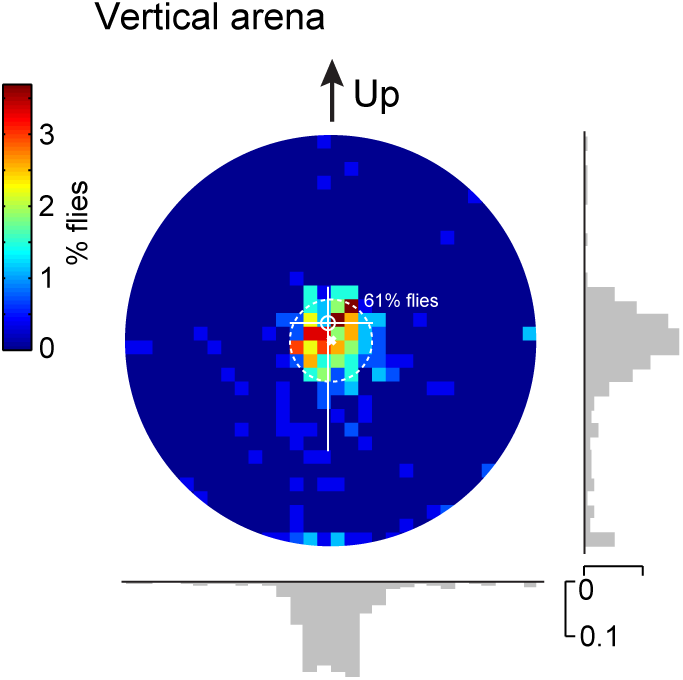
The shutter mechanism effectively relocates flies to the center of the arena, related to Figure 1. Most flies begin each trial in the center of the arena, as shown by the distribution of flies (vertical arena, start of a trial). The dotted white circle shows the minimum arena radius achieved during pre-trial shutter movements. After opening of the shutter, 61% of flies are still within this area. White circle indicates the mean starting position of all flies and the white lines indicate the 0.1 and 0.9 quantiles of the fly starting positions (in two directions). Two further effects of the shutter are visible: some flies are pulled outward along with the blades and some flies fall to the bottom of the arena if it is tilted. Same data as presented in Figure 1D, vertical arena.

**Figure S2.**
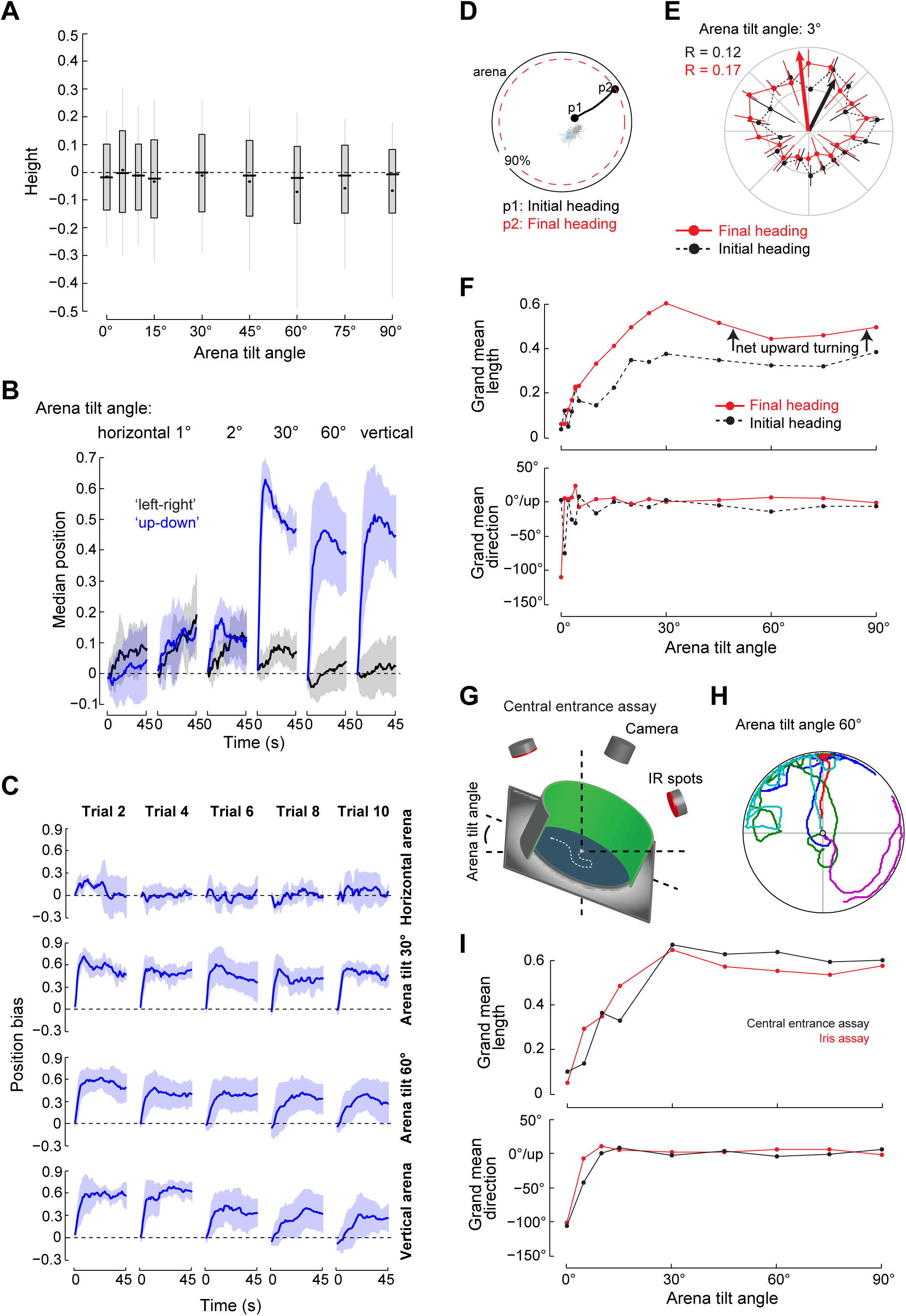
Gravity response at different arena tilt angles, related to Figure 2. (A) The distribution of fly start positions at the beginning of a trial shows that while higher arena tilt angles lead to more flies starting below the center of the arena, the majority of flies start a trial from the center of the arena. Standard boxplots, indicating 0.1, 0.25, 0.75 and 0.9 quantile ranges, black lines mark median, black dots mean start positions. N ≥ 4 experiments per tilt angle, see Figure 2. (B) A comparison of the median fly position over time along the tilt axis (up-down) and the perpendicular (left-right) axis shows that the arena does not impose a directional bias (mean ± s.d.). In the horizontal case, no directional preference is visible for either direction, while tilting the arena leads only to a preference for the ‘Up’ direction. Same N ≥ 4 experiments per tilt angle, see Figure 2. (C) The position bias averaged across experiments (mean ± s.d.), but shown for the indicated trials and arena tilt angles. At higher tilt angles there is a decrease in the amplitude of the behavioral responses that is not present at the lower tilt angles. Generally, the overall negative gravitaxis response remains, but this may be an effect of adaptation to the repeated startle. N ≥ 4 experiments per tilt angle, see Figure 2. (D) Initial fly responses, directly at the start of a trial were analyzed by comparing initial body orientations (p1) with their walking direction at crossing 90% arena radius (final heading, p2). For this analysis, we selected flies that started in the center of the arena (distance to edge >=0.5). (E) The initial heading at p1 (black) is compared with the final heading at p2 (red), for an arena title angle of 3°.. Arrows represent the corresponding grand mean vector directions. The polar plot shows the mean (± s.d.) number of flies in 20° bins. The increase in vector length indicates upward turning of flies, even at a tilt angle of 3°. (F) Grand mean vector lengths and directions are plotted for the initial fly heading distributions (black) and final fly heading distributions (red) across arena tilt angles. This indicates a net upward turn after the start of a trial. The Moore test shows that uniformity is significantly rejected for all tilt angles >= 3° (p=0.001) and not rejected for the horizontal arena (p>0.05). This confirms the low detection threshold found by using the gravitaxis scores. N ≥ 4 experiments per tilt angle, see Figure 2. (G) An additional behavioral assay was used to test the directional responses of individual flies. In this simpler assay, flies entered a tilted arena through a hole in the center. Arena diameter was 55 mm and the center hole (3 mm diameter) usually allowed 1 fly to emerge at a time. An experiment lasted 10 minutes. The experiments were conducted in the dark; the arena LEDs were not illuminated. Subsequent data processing was identical to the shutter assay. (H) Five example trajectories obtained in the central entrance assay with the arena tilted to 60*° are shown*. (I) *Initial fly responses were analyzed to compare both assays. For the iris assay, we used the analysis described above (D-F), but selected flies that begin with a distance to edge >= 0.75 (the flies that start very close to the center). Iris assay data* N ≥ 4 experiments per tilt angle, see Figure 2*; central entrance assay results based on 4 experiments per arena tilt (with 71-88 flies analyzed per arena tilt angle)*.

**Figure S3.**
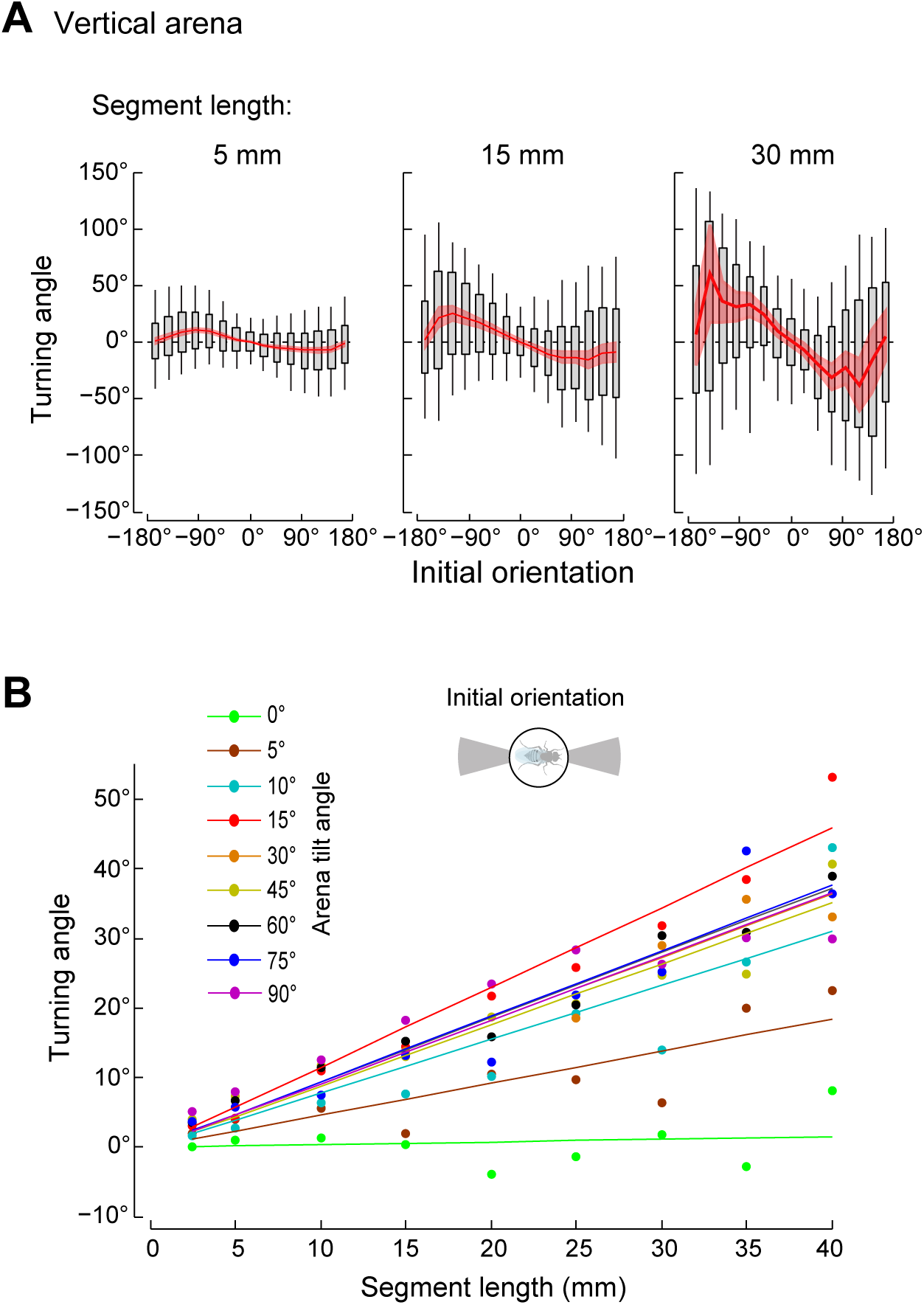
Fly turning analysis is robust to choice of trajectory segment length, related to Figure 3. (A) The turning angle analysis of Figure 3B was repeated for segment lengths of 5, 15, and 30 mm and plotted using standard boxplots, indicating 0.1, 0.25, 0.75 and 0.9 quantile ranges; red line represents the mean response vector direction, red area the 95% confidence interval. (B) The segment turning angle as a function of segment length. The turning behavior was analyzed by pooling trajectory segments with an initial sideways orientation and then examining the turning response for different arena angles. For arena tilt angles above 5°, the turning response was monotonically (and approximately linearly) related to the segment length used for analysis, indicating that flies continue to reorient upwards over several length scales.

**Figure S4.**
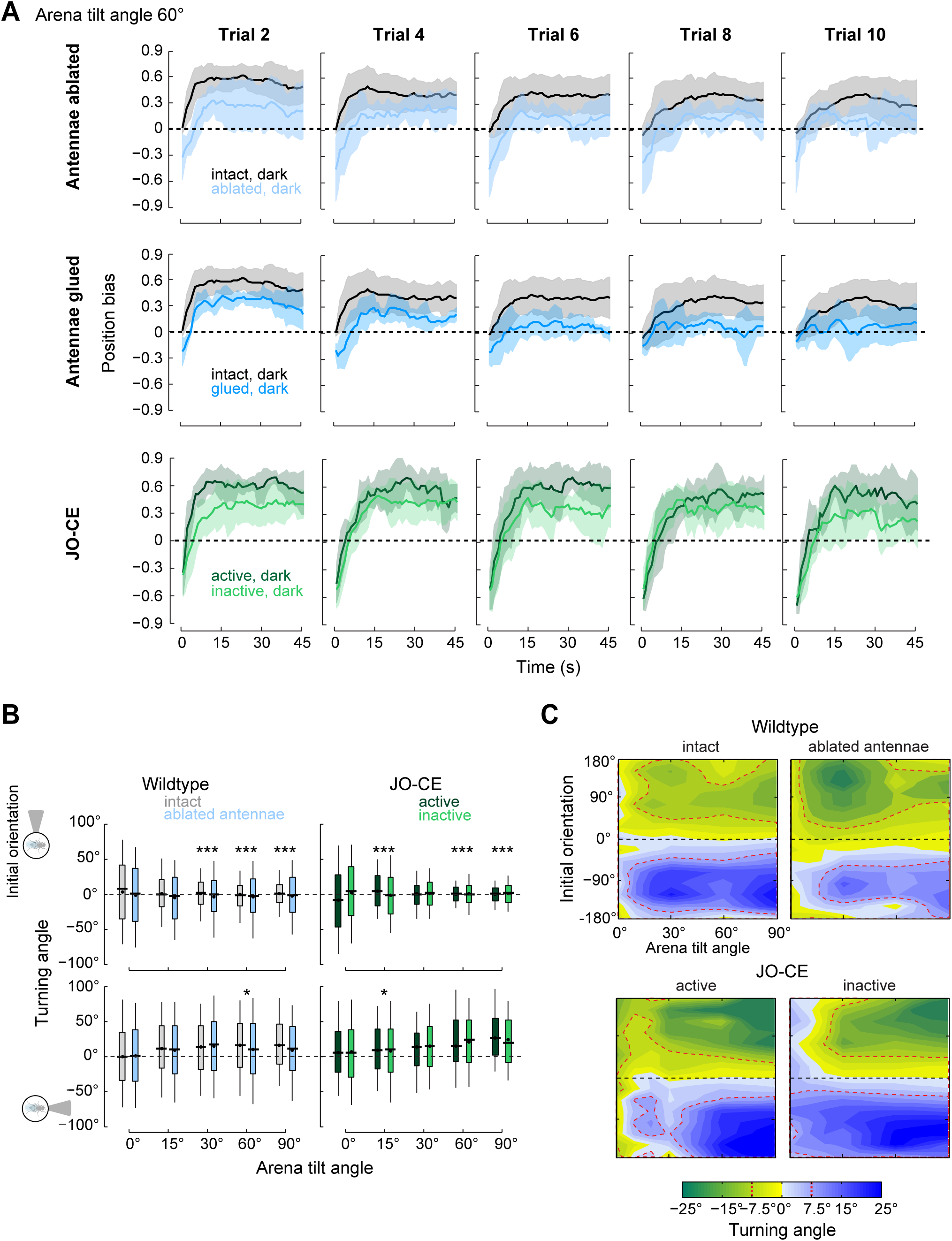
Influence of antennal ablations on orientation behavior, related to Figure 4. (A) The effects of manual and genetic antennal manipulations are compared across trials for an arena tilt of 60*°. In each case, the controls (black/ dark green) are compared with the effects of manipulations (blue/light green). This shows that adaptation effects between controls and antennal manipulations were similar and did not influence the general observation that antennal manipulations led to an overall lower median position bias. (based on data presented in* Figure 4, N ≥ 4 experiments / condition). (B) *The variance of turning behavior is compared for surgical and genetic ablation of the antennae, for flies that are initially oriented upwards or sideways. For each tested arena angle, we compared the variance of the segment turning angles with an F-test (based on data presented in* Figure 4, N ≥ 4 experiments / condition). (C) The turning behavior depends on both fly orientation and the arena tilt angle, and is shown for manual and genetic antennae manipulations (and control conditions). To aid in the visualization of these data, we use a turn angle threshold of 7.5° (red dotted lines). The black dashed line shows the upward direction. *Results based on experiments and arena tilt angles presented in* Figure 4*, using same analysis method as in* Figure 3C.

**Movie S1. *Drosophila* walk upward in a tilted arena, related to Figure 1.**

A comparison of a typical trial of flies walking in a horizontal arena and the same arena inclined by 60 degree. The different phases of the experiment (see Figure 1B) are marked with green lines in the lower plot. At first, flies are pushed to the arena center, then they are mechanically startled, the arena edges are opened, and then trial commences. At the start of the trial, the fly tracking results are superimposed on the recorded videos. Triangles show the position of individual flies and their orientation. White lines on the video data show the 0.1 and 0.9 quintiles of fly position; the cross indicates the overall mean fly position. Colored dots represent the median position bias of flies in each arena. The median position bias is shown over time in the lower plot.

**Movie S2. The antennae are dispensable for gravitaxis, related to Figure 4.**

The gravitaxis behavior of intact wildtype flies is compared against wildtype flies with bilateral antennae ablations. Experiments were performed in an arena tilted to 60 degree. The fly positions obtained by tracking are superimposed on the recorded video. White lines on the video data show the 0.1 and 0.9 quantile range of fly position; they cross at the mean fly position; black squares on the vertical axis indicate the median fly position. The lower analysis plots show the median start position along the vertical axis and the number of flies falling during the pre-trial startle and during each trial. During the iris opening sequence before each trial, the video is slowed to 50% speed to better illustrate the impact on flies. Green represents intact wildtype results, magenta antennae ablated wildtype flies. For the purposes of this movie only, these trials are 20 seconds long.

## Materials and Methods

### Experimental animals

*Drosophila melanogaster* flies were reared on standard media at 25 °C on a 16/8 hour light/dark cycle. Flies were cold-anesthetized (∼4 °C), sorted, and placed into vials containing standard media and then used for experiments within 4 hours. Directly preceding the experiment, flies were again cold-anesthetized and placed in the arena. Experiments were started after a recovery time of approximately 10 minutes, a point in time at which most flies actively explore the arena.

Our standard experimental animals were 3-5 day old wildtype *Drosophila melanogaster* (of the so-called DL strain, derived from wild-caught flies and generated in Michael Dickinson’s laboratory around 1995). We used mixed populations of both males and females, since initial tests did not reveal a sex effect on gravity behaviors. Neural silencing experiments of antennal Johnston’s organ neurons were performed with NP6250, a GAL4 driver line that targets subgroups C/E of the JO neurons [1]. NP6250 was obtained from the Bloomington *Drosophila* stock center. This line was crossed to w+(DL);UAS-cTNTe,tubP-Gal80ts;+, obtained from Tyler Ofstad (which was backrossed for 10 generations into the DL background to control for background effects [2]). Flies were reared at 18° and two day old adult flies were then temperature shifted to 30° for 40-48 h prior to the experiments to express tetanus toxin in the target neurons. Flies without the temperature shift, but of the same age range, were used as controls. All experiments were performed at our room temperature (22°-24°).

Manual manipulations of sensory structures of DL flies were performed under cold-anesthetization. Ablations of antennal segments and halteres were performed with fine forceps, 24-28 hours prior to the experiments. Wings were ablated with fine scissors 1-5 hours before the experiments. Fixations of antennal joints or the head-thorax joint (neck) were performed with UV light-curing glue, 1-5 hours before the experiments.

### Behavioral experiments

To examine the influence of gravity on fly locomotion, we used an arena similar in design to Ofstad et al. [2], and adapted it to a tilting platform. The arena had a diameter of 117 mm and a height of about 3 mm. The top of the arena was covered with a glass disk, whose inner surface had been coated with a slippery silicone film (Sigmacote, Sigma-Aldrich, St. Louis, USA). The 3 mm arena height enables normal fly locomotion while preventing flight; the slippery coating prevents flies from walking on the lid. The arena was placed inside of a dark enclosure within a room that was lit using low intensity red lighting. This should prevent nearly all visual cues. The arena itself was illuminated with infrared lights (Smart Vision Lights, Muskegon, USA) and fly behavior was recorded with a CMOS camera at 15 frames per second, fitted with an infrared filter (Basler 601f, Basler AG, Ahrensburg, CH). Approximately 10 minutes prior to an experiment, flies were placed in the arena and the tilt angle was set between horizontal (0°) and vertical (0°). Exact tilt angles were calibrated with an inclinometer to sub-degree resolution. After an experiment, the arena was cleaned with 70% ethanol.

Walking flies exhibit spatial preference for vertical objects and edges (Figure 1D, Distance to edge) [3, 4]. As our analysis required flies to freely orient with regards to the arena slope, we developed a mechanism that allowed arena radius changes to relocate flies to the center of the arena for repeated testing (Figure 1B). In principle, this mechanism worked like a camera iris, although at much lower speeds. Using a design of interleaved blades that could be rotated into each other, we achieved radius changes between 15 mm and 63.5 mm with variable, computer-controlled speed profiles. Interleaved blades allowed us to achieve the necessary arena height while preventing flies from crawling between the blades. The iris movements are controlled through an RC servo (HS-805BB, ServoCity). Before each trial, a sequence of radius changes was used to (I) slowly push flies to the center of the arena, (II) provide a mechanical startle, and (III) quickly open the arena to start the trial (Figure 1B).

Different lighting conditions were created by placing a single blue LED (470 nm peak) at the perimeter of the arena for phototaxis stimulation (Figure 2D) or placing a circular LED array on top of the arena (Figure 2E) to provide uniform, green illumination (568 nm peak) at different intensities [5]. The experimental procedures and conditions were controlled through Matlab (The MathWorks, Natick, MA) and for each experiment, video data (uncompressed AVIs, one per trial) and associated experimental metadata were saved for offline analysis.

### Data Processing

Fly trajectories were obtained from the video data using the Ctrax fly tracking software [6], resulting in body position and body orientation time series. It was not possible to obtain fly identities across trials since we did not track the flies during the arena closing and opening sequence. Position data were smoothed with a 1/3 s window moving average (to facilitate speed measurement, Figure 1D). Metadata and trajectory time series were stored in a custom MySQL (Oracle Corporation, Santa Clara, USA) database for further analysis. Data were selected from the database and retrieved into Matlab for analysis and visualization (Mathworks, Natick, USA).

### Behavioral metrics and statistics

To analyze the gravity-specific behavioral responses, we primarily used the *position bias* of the flies, which is the distribution of the ‘vertical’ height of the flies relative to the boundaries of the arena (Figures 1D; 2C; S2C and S4A). Further behavioral metrics such as the speed or the distance to the arena edge were used (Figure 1D). Distance to the arena edge is an absolute, normalized value where 1 indicates a fly is in the center of the arena and 0 means that a fly is at the edge (Figure 1D). To better compare across experiments and conditions, we defined the *gravitaxis score* to be the median position bias at 30 seconds (Figures 1D; 2A, B, D, E; 4A, B). Unless otherwise indicated, these metrics are determined for the data pooled across the 10 trails. For each experimental condition and genotype, we then calculated the mean gravitaxis score over experiments. A two-tailed T-Test was used for statistical comparison of gravitaxis scores across experimental conditions and genotypes. Throughout the manuscript we use the same convention for assigning significance to figures (* p<0.05, ** p< 0.01, *** p < 0.001).

Fly turning behavior was analyzed by pooling data across experiment (with identical conditions) and then dividing all fly trajectories into non-overlapping segments (Figures 3; 4D; S3; S4B,C). In Figures 3 and 4, these segments are 15 mm in length, but the different segment lengths are explored in Figure S3. As turning behavior of flies walking along the edge of the arena is restricted and flies like to follow walls, we excluded segments close to the arena edge (segments that exceeded 90% maximum arena radius). For each segment, we calculated a turning angle as the difference between the fly body orientation at the end and the beginning of the segment (we also did this analysis based on trajectory orientation with similar results, not shown). For further comparison of segment turning we binned initial segment orientations (24° bins) and analyzed the distribution of segment turning angles for each bin. Using circular statistics for each bin (of the initial segment orientation), we calculated the mean response vector and the corresponding 95% confidence interval [7]. The direction of the mean response vector was used as an indication of the mean segment turning angle (Figures 3B; 4D; S3A, B).

An F-Test was used to compare the distribution variances of turning angles within individual bins across arena tilt angles and conditions (Figures 3D; S4B). A Mann-Whitney U-Test was used to compare turning response amplitudes of individual bins across arena tilt angles and conditions (Figure 3D).

Fly turning behavior was compared across arena tilt angles by plotting the mean segment turning angles as a contour plot. For this plot, we interpolated the previously obtained mean segment turning angles for each arena tilt angle and initial segment orientation. 10° steps were used for interpolating data along the arena tilt angle axis and 24° for interpolating along the initial segment orientation axis. Finally, mean segment turning angles were plotted using false color (as indicated by the color code; Figure 3C and S4C).

In Figure S2D-I, we analyze the initial responses of flies. For this analysis, only flies that started in the center of the arena at the beginning of a trial were used (Distance to edge >=0.5). We examined the initial orientation of these flies (at the start of each trajectory) and also their orientation when crossing the circle corresponding to 90% of the maximum arena radius (which we referred to as ‘final heading’). For each arena tilt angle we calculated the grand mean vector (using circular statistics) for the initial and final orientations across experiments [7]. The direction of the grand mean vector represents the average initial or final walking direction, while the length of the grand mean vector is a measure of the variance in directedness. The Moore test was used for statistical comparison of grand mean vectors (see legend Figure S2F) [7].

A simpler behavioral setup (the central entrance assay, Figure S2G-H) was also used to test the directional responses of individual flies. This setup was not outfitted with the iris mechanism, and individual flies would enter a tilted arena through a hole in the center. Arena diameter was 55 mm and the center hole a diameter of 3 mm, usually allowing 1 fly to emerge at a time. An experiment lasted 10 minutes. Subsequent data processing was identical to the standard iris assay.

